# Distances walked by beach users and protecting shorebird habitat zones from disturbance

**DOI:** 10.1101/783696

**Authors:** Stephen Totterman

## Abstract

Human population growth along Australia’s coast is increasing development and recreation pressures on beaches and shorebirds. This study observed human recreation on 18 beaches on the far north coast of New South Wales in February and March 2019. The far north coast supports the largest numbers of beach-resident Australian Pied Oystercatchers *Haematopus longirostris* in the State. The most frequent activities observed were walking (29%), dog walking (21%) and swimming (16%). Walkers covered greater distances compared to other beach users. For beach walkers and dog walkers combined, the mean along shore distance walked from an access point was 809 m and the 95th percentile was 1990 m. Noting that human recreation disturbance is a major conservation threat to beach nesting birds and that pro-environmental behaviour is uncommon among beach users, large separation distances between beach access points and shorebird habitat zones are recommended to reduce human intrusions into those zones. This spatial zoning and passive human exclusion strategy can be applied to long and less-developed beaches.

## INTRODUCTION

Growing human populations and demand for coastal lifestyles away from the major cities (the “sea change” phenomenon) are strong demographic forces that are increasing development and recreation pressures on Australia’s coastal areas (Priest *et al.* 2002). BirdLife Australia (2018) have stated that the greatest conservation threat to beach nesting birds is disturbance from people visiting the beach.

Birds typically respond to a perceived threat, such as a potential predator or a human, by moving away (Blumstein 2006). Frequent human recreation disturbance can result in displacement of shorebirds from their preferred habitat (Schultz & Stock 1993, Liley & Sutherland 2007), increased energy expenditure in flight responses (West *et al.* 2002), reduced energy intake through disrupted foraging (Burger 1994, West *et al.* 2002) and exposure of eggs or dependent young to diminished parental care (Lord *et al.* 2001, Verhulst *et al.* 2001, Harrison 2009). Disturbance has the potential to result in declines in bird populations if displacement becomes permanent and there is little surplus habitat to move into, if survival is reduced or if breeding success is lowered (Gill *et al.* 2001). However, it is difficult to extrapolate observed individual or local effects to population impacts (Davidson & Rothwell 1993, Hill *et al.* 1997, Gill *et al.* 2001).

Human recreation disturbance rates on beaches often outnumber natural disturbances, even on beaches with low visitation rates (Weston & Elgar 2005, Harrison 2009). Habitat and life history attributes make beach nesting birds particularly sensitive to human recreation disturbance. Ocean beaches are narrow, linear habitats where birds can find it difficult to maintain a safe distance from people (Pienkowski 1993, Harrison 2009). Territorial behaviour and high site fidelity of oystercatchers (Taylor *et al.* 2014), for example, means that breeding pairs are unlikely to relocate to alternative sites, if they exist. Australian pied oystercatchers are large shorebirds (length 42–50 cm; Marchant & Higgins 1990) and large birds are generally more “flighty” than are smaller species (Blumstein 2006, Weston *et al.* 2012a). Beach nesting bird populations are often concentrated at a number of sites and local changes in breeding populations at those sites are population impacts.

Manfredo and Dayer (2004) noted that the thoughts and actions of humans ultimately determine the course and resolution (if any) of human-wildlife conflicts. Practical research into human recreation disturbance of wildlife must then include studies of human attitudes and behaviour (*e.g.* Antos *et al.* 2007, Maguire *et al.* 2011, Guiness *et al.* 2020, Schneider *et al.* 2020). However, far more studies into human disturbance of birds have simply measured flight initiation distances (FIDs), *i.e.* the distance at which an individual bird or flock moves away from a stimulus (typically a human walking towards the focal bird or flock) (see Weston *et al.* 2012a for a summary). A common application for FID data is for selecting “buffer distances” between humans and wildlife. However, Hill *et al.* (1997) cautioned that there are potentially many experimental variables that affect FIDs (species identity, individuals or flocks, breeders or non-breeders, the habitat, activity of the birds prior to disturbance, stimulus applied, starting distance, *etc.*) and that the results of any one study are often limited to specific experimental conditions. There is a risk that end users may generalise and misuse FIDs when developing management plans for human-wildlife conflicts.

Codes of conduct and enforceable regulations are commonly applied with the aim to reduce human-wildlife conflict. However, evidence is growing that “pro-environmental” behaviour is uncommon in coastal areas and that compliance with regulations is low (Dowling & Weston 1999, Antos *et al.* 2007, Weston *et al.* 2014, Maguire *et al.* 2019, Schneider *et al.* 2020). Glover *et al.* (2011) surveyed 295 visitors and residents at Western Port Bay, Victoria and reported that while 86% of respondents supported protection of shorebird habitat, only 39% were supportive of limitations on access for walkers to avoid disturbance to wildlife. Given that it is difficult and costly to manage human behaviour, the application of FID data for shorebirds on beaches may be limited to designing human exclosures around individual nest sites or colonies and shorebird roosts (*e.g.* Lafferty *et al.* 2006, Weston *et al.* 2012b).

This study recommends, for longer and less-developed beaches, spatial zoning to restrict disturbance activity to certain areas of the beach (Hill *et al.* 1997). Large separation distances between access points and shorebird habitat zones (*i.e.* greater than the distances people typically walk) can reduce intrusion rates into those zones and disturbance to shorebirds (Dowling & Weston 1999, Antos *et al.* 2007, Liley & Sutherland 2007).

Selecting an appropriate separation distance depends on visitation rates to the beach. For the simplest case of one habitat zone and one access point, the intrusion rate (*R*) into the habitat zone (*e.g.* persons per hour) is:

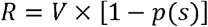

where *V* is the visitation rate, *s* is the separation distance between the access point and the start of the habitat zone and *p*(*s*) is the proportion of beach users walking to the start of the zone (Figure 1).

**Figure 1.**
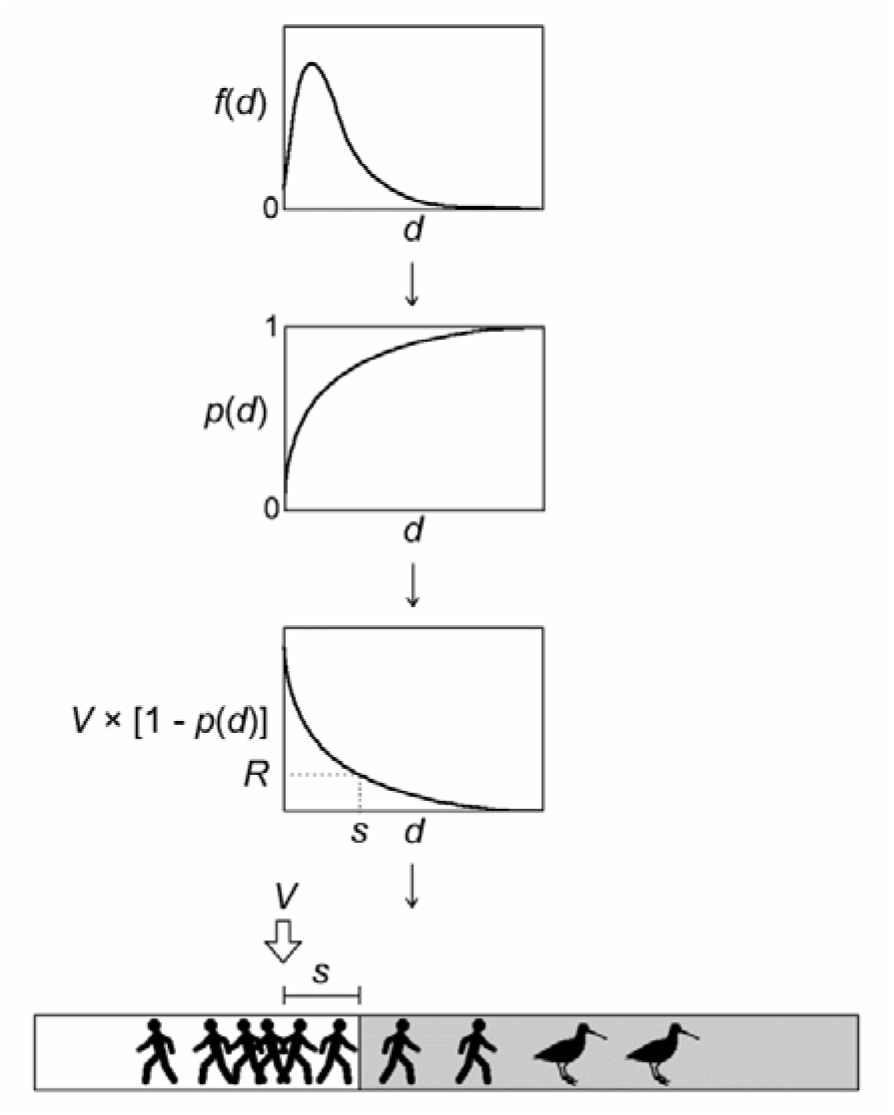
Beach user intrusions from an access point (arrowed) into a shorebird habitat zone (grey area). The distributions are (from top to bottom) a frequency distribution of distances walked (*f*(*d*)), a cumulative frequency distribution (*p*(*d*)) and a distribution of intrusions (*V* × [1 – *p*(*d*)]). *V* is the number of people using the access point and walking along shore towards the habitat zone. The separation distance is *s* and *R* is the number of intrusions into the habitat zone.

Given a fixed intrusion rate (*e.g.* potential disturbance events per hour), the separation distance must be increased as the visitation rate *V* increases. Recognising strong among person/group variability in distances walked and anticipating future growth in human populations and beach recreation, it is recommended to simply adopt the 95th percentile distance under the assumption that patterns of beach recreation and distances walked hold into the future. However, as daily visitation rates increase to hundreds of people or more, other management actions that attempt to promote coexistence between humans and shorebirds could be investigated (reviewed by Pienkowski 1993).

This study observed distances walked for beach users on the far north coast of NSW. This region supports the largest numbers of beach-resident Australian Pied Oystercatchers *Haematopus longirostris* in the State including the South Ballina Beach, Broadwater Beach (which includes Airforce Beach and Salty Lagoon), Bombing Range Beach and Yuraygir Key Management Sites currently recognised by the NSW Department of Planning, Industry and Environment (2019). The Australian Pied Oystercatcher is currently listed as Endangered in Schedule 1 of the NSW Biodiversity Conservation Act 2016 (NSW Government 2016).

Preceding studies for the Australian Pied Oystercatcher have reported mean flight initiation distances from 39–129 m (Table 1). This study was motivated by concerns that oystercatchers at South Ballina are increasingly threatened by “beach house” developments and that misuse of FID data has resulted in inadequate separation distances between new beach accesses and oystercatcher territories. For example, the FID results of Harrison (2009, Table 1) could be misused to allow access points as close as 46 m to oystercatcher nests. There are several problematic assumptions underlying this choice, including: nest locations are fixed and do not change from year to year, a breeding pair needs only 0.7 Ha of space (*i.e.* a circular area with radius 46 m) and people will not bring domestic dogs onto the beach, *etc*.

**Table 1.** Mean flight initiation distances (FIDs) for Australian Pied Oystercatchers in preceding studies. For comparison, the mean distance walked by beach users in this study was 444 m (see Results).

| Reference | Habitat | Stimulus | Mean<br>FID (m) | Breeding<br>status |
| --- | --- | --- | --- | --- |
| Paton *et al.* 2000 | Estuary | Human | 83 | Non-breeding |
| Blumstein 2006 | Unknown | Human | 39 ($n = 23$) | Unknown |
| Harrison 2009 | Beach | Human | 46 ($n = 79$) | Breeding |
| | | Human and dog | 129 ($n = 20$) | Breeding |
| Glover *et al.* 2011 | Varied | Human | 42 ($n = 21$) | Non-breeding |

## METHODS

### Study sites

Twenty locations (sites) were sampled on 18 beaches between Tweed Heads and Coffs Harbour, NSW, between 8 February and 24 March 2019 (Figure 2, Table 2). Five oystercatcher breeding beaches were included. Most beaches were moderate to long (mean length 6.5 km). On shorter beaches, *e.g.* Chinaman’s Beach (length 0.8 km), walking distances are constrained by the beach length.

**Figure 2.**
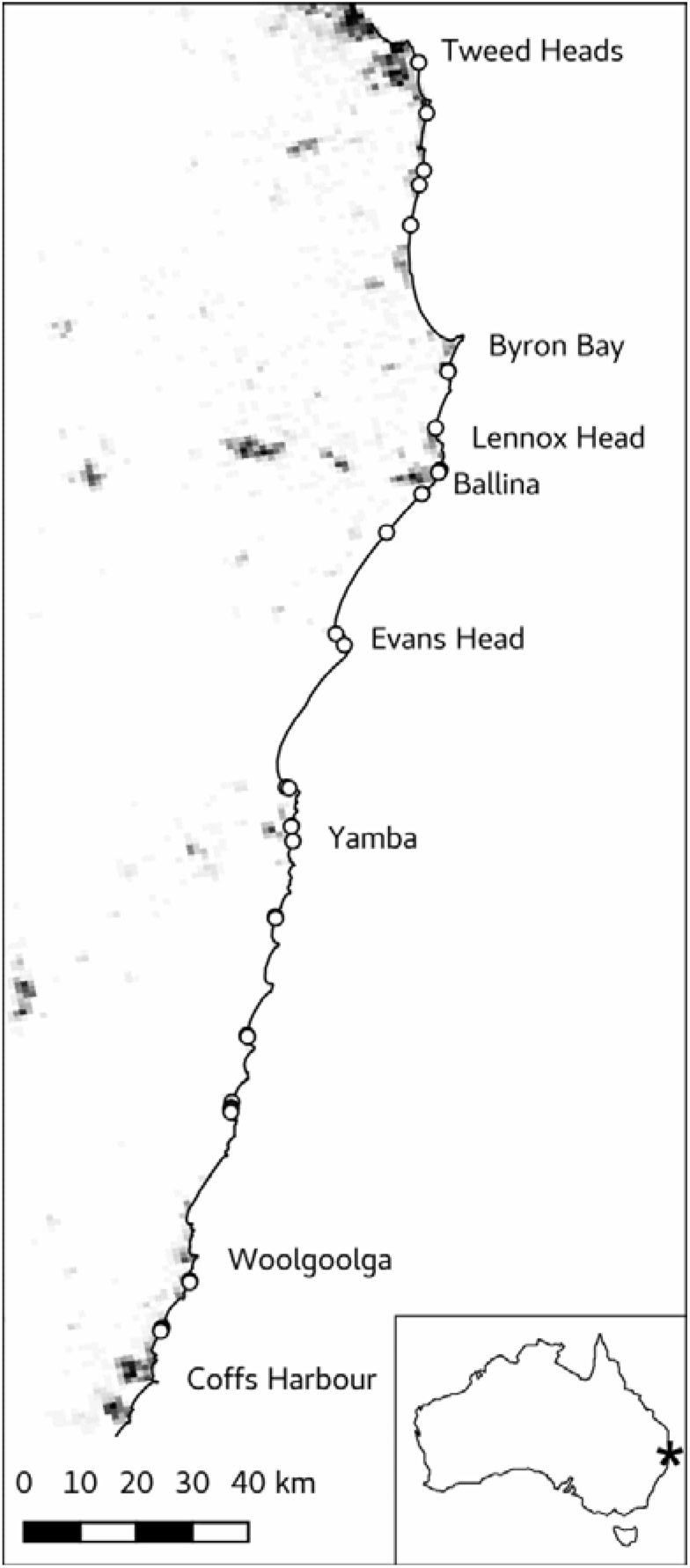
Map of the far north coast of New South Wales showing the location of access points (white-filled circles) where beach recreation was studied (see Table 2 for details of beaches and sites). The background image shows human population densities (scale 0–2000 persons.km^−2^; darker values indicate higher densities) from the Australian Population Grid 2011 (https://www.abs.gov.au/ausstats/abs@.nsf/mf/1270.0.55.007). Text annotations identify larger coastal towns and cities. The inset map shows the location of the study region within Australia.

**Table 2.**
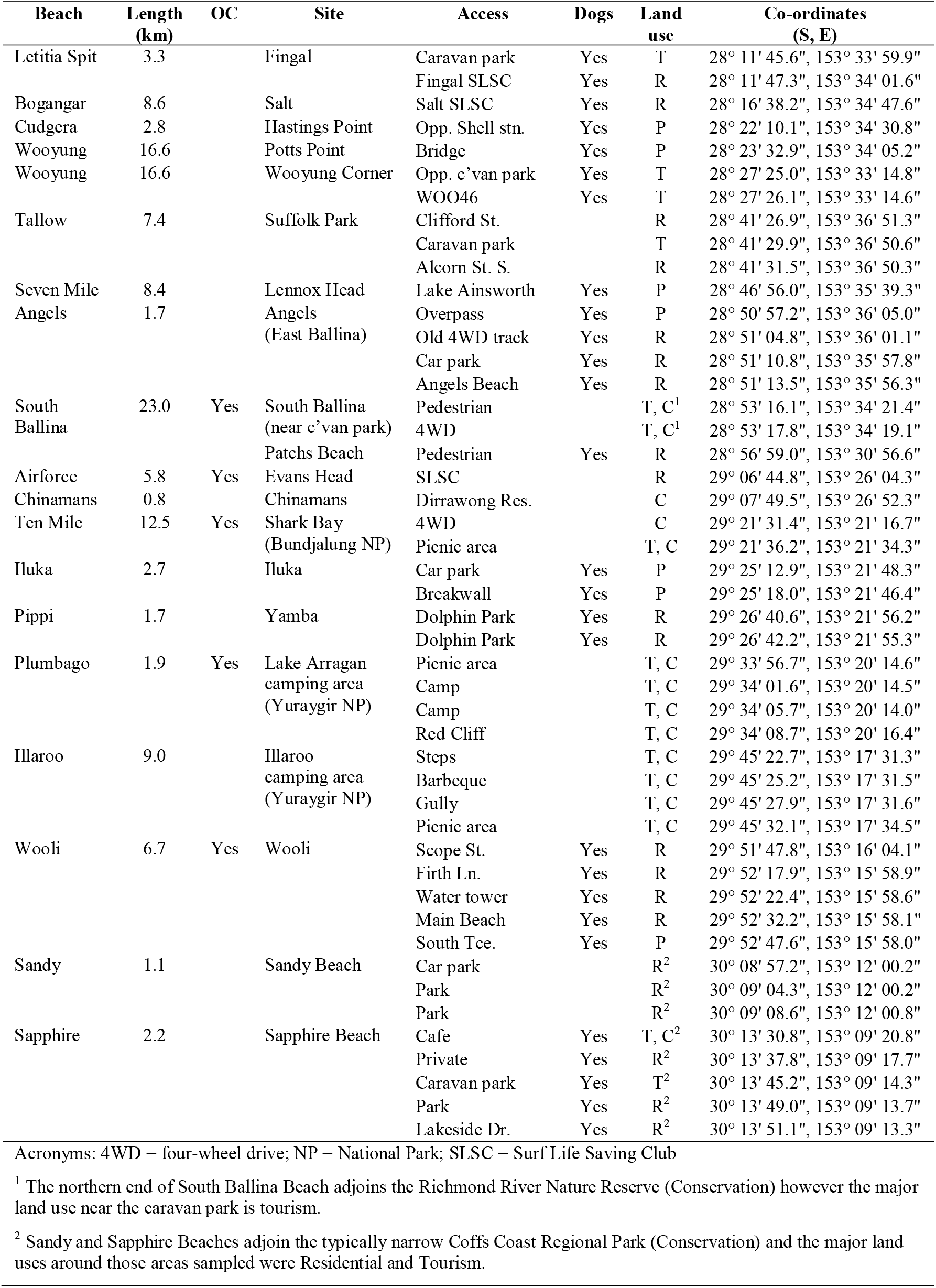
Study sites and beach access points sampled in this study (north to south). Beaches known to have breeding oystercatchers (OC) are recorded as ‘Yes’. Access points where dogs are allowed onto the beach are recorded as ‘Yes’. Major land uses within 200–300 m of each access point are classified into four categories: R = Residential; P = Public Reserve (typically at ‘peri-urban’ sites); T = Tourism; C = Conservation.

Sites were selected to include a variety of adjoining land uses and dogs permitted and dogs prohibited zones (Table 2). Thirteen sites had a number of closely spaced access points that were observed together as one site. The mean between-site distance of 87.7 km was far greater than the mean within-site distance of 353 m (for sites with multiple access points). Remote and infrequently used sites are not productive and were not sampled. For example, visitation rates to the picnic area north of Broadwater Headland (at the southern end of South Ballina Beach) were too low to persevere with sampling.

### Observations

Sampling occurred in the morning and on any day of the week. There were no school holidays during this study. Ten observations of beach users were made at each site. Social groups of beach users (*e.g.* a picnic) were recorded as one observation, to avoid pseudoreplication. The total sample size was 200 observations (20 sites × 10 observations per site).

For busy access points, every third person/group entering the beach was observed, counting from when the observer was within sight of the access point and ready to record new observations. For less busy beaches or at less busy times, accumulation of observations was accelerated by observing every consecutive person/group and/or observing multiple access points. Patchs Beach was visited twice to accumulate ten observations and South Ballina was visited six times. Sampling of other sites was completed in one visit.

The response variable was the maximum one-way distance a focal person/group walked from an access point. Distances were measured using a handheld Global Positioning System (GPS) satellite receiver (Garmin GPSmap 62s) or estimated if less than *c.* 50 m (where GPS imprecision is large relative to the distance to be measured). Distances were measured in the along shore direction and included zero distances where the person/group did not move along shore from the access point (*e.g.* walking directly down shore to the water). Focal persons/groups were observed discreetly from some distance to avoid disturbing them and potentially affecting their behaviour. Binoculars were used as a visual aid and, on longer beaches, a bicycle was used (at low tide) to quickly move between observation points.

Predictor variables recorded together with each focal person/group observation were: day, time the person/group entered the beach (Australian Eastern Standard Time, without daylight saving), number of adults in the group, number of children in the group, numerically dominant gender of adults in the group (male, female or balanced), number of accompanying dogs, activity of the person/group, count of other people within 500 m of the access (excluding the observer), presence/absence of lifeguards on the beach, nearest tide (high or low), a visual estimate of percent cloud cover, wind speed in five subjective categories (calm, very light, light, moderate, strong) and wind direction (eight cardinal points: N, NE, E, SE *etc*). Daily 9 am temperature (°C) and rainfall (mm) observations from the nearest Australian Bureau of Meteorology weather station (http://www.bom.gov.au/climate/data/index.shtml) were later added to the data table. Other variables recorded were access type (pedestrian, vehicle, combined) and distance to parking (m) for each access point and beach length (km) for each site.

Group composition variables (adults, children, gender, *etc.*) were recorded because they may affect the motivation or capability for walking long distances. Counts of people on the beach were recorded because new arrivals may have to walk some distance to find their own space. Similarly, the presence/absence of lifeguards was recorded because people may have to walk some distance to swim in the patrolled zone. Tide was recorded because the firm intertidal zone sand at low tide can facilitate walking longer distances. Weather variables were recorded because uncomfortable conditions can deter walking long distances. Time of day was recorded because it is related to weather conditions and because some persons would have been walking early before work and had limited time. Weekday was recorded because some persons may have had more time for recreation on weekends. Distance to parking was recorded because it could affect the motivation or capability for additional walking along the beach. Access type was disregarded because there was little variation for this variable (96% pedestrian access points).

All 10 activities recorded are described in Table 3. Only activities that involved walking onto and along the beach were recorded. Joggers, runners, cyclists and horse riders were not included in this study because, even though they often cover longer distances than walkers, those activities were uncommon. Four-wheel drives (4WDs) were not part of this study and beach driving was permitted on only seven of the 18 study beaches.

**Table 3.** Descriptions of beach recreation activities recorded in this study.

| **Activity** | **Description** |
| --- | --- |
| Walking (WK) | Walking along the beach, usually direct and close to the shoreline. Walkers walk for some distance and then turn back or exit the beach via a different access point. |
| Dog walking (WD) | Walking along the beach accompanied by one or more dogs, on or off leash. |
| Swimming (SW) | Does not include wading (to approximately knee-depth). |
| Surfing (SF) | Mostly on surfboards. |
| Picnic (PC) | Picnics generally involved multiple persons and a variety of activities. Features of this activity can include beach shelters, food and beverages, recreational equipment and the group stays in one location for an extended period. |
| Sunbathing (SB) | Self-explanatory. |
| Looking (LK) | Viewing the beach or ocean, checking the beach, fishing or surf conditions and often where there is seating on the beachfront. |
| Fishing (FI) | Recreational line fishing. |
| Playing (PL) | Young children playing in the sand. Unlike picnickers, these groups don't bring so much equipment, don't stay long and adults present are supervising and not engaged in other recreation. |
| Photography (PH) | Recreational photography. |

### Statistical analysis

Conditional inference regression trees (Hothorn *et al.* 2006) were used to discover relationships between distance walked and predictor variables. Regression trees are a machine learning method that explains variation in a single variable by repeatedly splitting the data into homogeneous groups using combinations of predictor variables (De’ath & Fabricius 2000). Machine learning methods can accommodate high-dimensional data, with many predictor variables (Breiman 2001). Conditional inference trees utilise statistical tests to avoid variable selection bias and overfitting.

Evaluation of machine learning models concerns the ability of models to predict new or withheld data rather than goodness of fit for a single model fitted to the all the data. A “useful” model is one that can make acceptably accurate predictions for new data. Accuracy (square Root Mean Squared Error; *RMSE*) of distance walked predictions was evaluated using five-fold cross-validation with ten repeats. Relative performance of regression tree models was compared to a mean distance walked null model and is reported as the reduction in cross-validated deviance: *D*^*2*^ = *(MSE*_*null*_ − *MSE*_*fitted*_*) / MSE*_*null*_ (Guisan & Zimmerman 2000).

The statistical analysis was performed in R version 3.5.2 (R Core Team 2018) and using the R packages *party* version 1.3-1 (Hothorn *et al.* 2018) and *caret* version 6.0-81 (Kuhn *et al.* 2018). Multiplicity adjusted *P*-values for conditional inference trees were computed using Monte Carlo permutation tests (with 9999 resamples), the minimum number of observations in a terminal node (‘leaf’) was set to 10 (5% of the sample size *n* = 200) and the minimum sum of numbers of observations in a node in order to be considered for splitting was set to 30 (3 × 10).

## RESULTS

A total of 200 observations of beach users were made from 20 sites on 18 beaches between 8 February and 24 March 2019. Twelve of the 20 sites (60%) were adjacent to residential areas and the other eight (Hastings Point, Potts Point, Wooyung Corner, South Ballina, Chinamans Beach, Shark Bay, Lake Arragan and Illaroo) were non-urban sites. Observations were nearly equally distributed between weekdays (51%) and weekends (49%) and high (43%) and low tides (57%). Most observations were early in the morning, between 0645–0930h AEST (interquartile range). Adults were present in 98% of observations and the mean group size was 1.4 adults (*n* = 197; 95% CI = 1.3–1.5). The dataset was nearly uniformly distributed among majority female (32%), male (38%) and balanced gender (30%) groups. The study took place at the end of a dry summer (mean rainfall 2 mm on sampling days). Mean 9 am temperature was 25 °C, mean cloud cover was 30% and wind speed was most frequently light.

The most frequent activities observed were walking (29%), dog walking (21%) and swimming (16%). The distribution of distances people walked had a strong positive skew (Figure 3). The mean distance was 444 m (*n* = 200; 95% CI = 365–523 m) and the 95th percentile distance was 1683 m, meaning that five per cent of persons/groups walked further than that distance. Walkers covered the longest distances compared to other beach users (Figure 4). Mean distances walked were similar for walkers with or without dogs at 748 m (*n* = 41; 95% CI = 605–892 m) and 851 m (*n* = 58; 95% CI = 667– 1035 m) respectively. For both walking activities combined, the mean distance was 809 m (*n* = 99; 95% CI = 687–930 m) and the 95th percentile distance was 1990 m.

**Figure 3.**
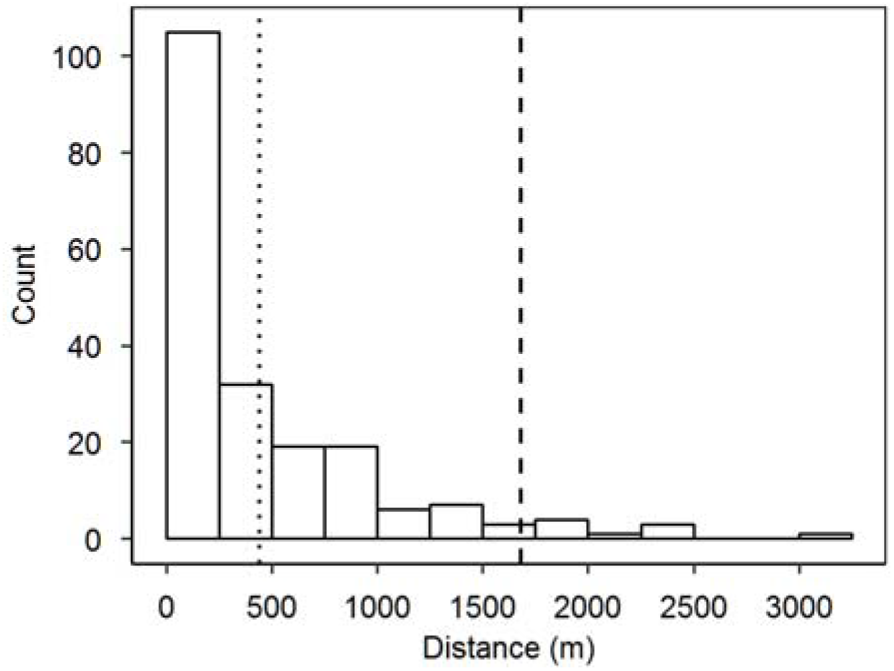
Distances walked for all beach users (*n* =200). Vertical dotted lines indicate the mean and 95th percentile distances (dotted and dashed lines respectively).

**Figure 4.**
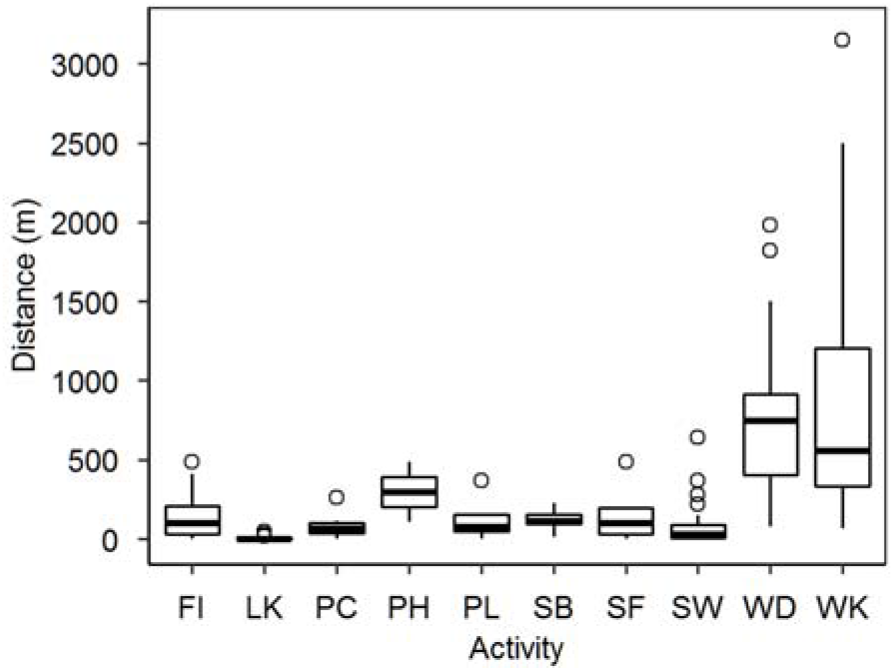
Distances walked for each activity: FI = fishing, LK = looking, PC = picnic, PH = photography, PL = playing, SB = sunbathing, SF = surfing, SW = swimming, WD = dog walking, WK = walking. Boxes show the first quartile, median and third quartile, whiskers extend to a maximum of 1.5 times the interquartile range and data outside the whiskers are plotted as individual points (white-filled circles).

A conditional inference regression tree model with minimum criterion 0.95 (1 − *P*-value) indicated that longer distances were associated with beach walking, with or without dogs (*P* < 0.001) and low wind speed (calm or very light winds; *P* = 0.048) (Figure 5). Other beach users walked shorter distances with a slight increase in the middle of the week (Wednesday and Thursday; *P* = 0.02), however there were only 17 observations in that node and that result is not expected to appear in replicate studies.

**Figure 5.**
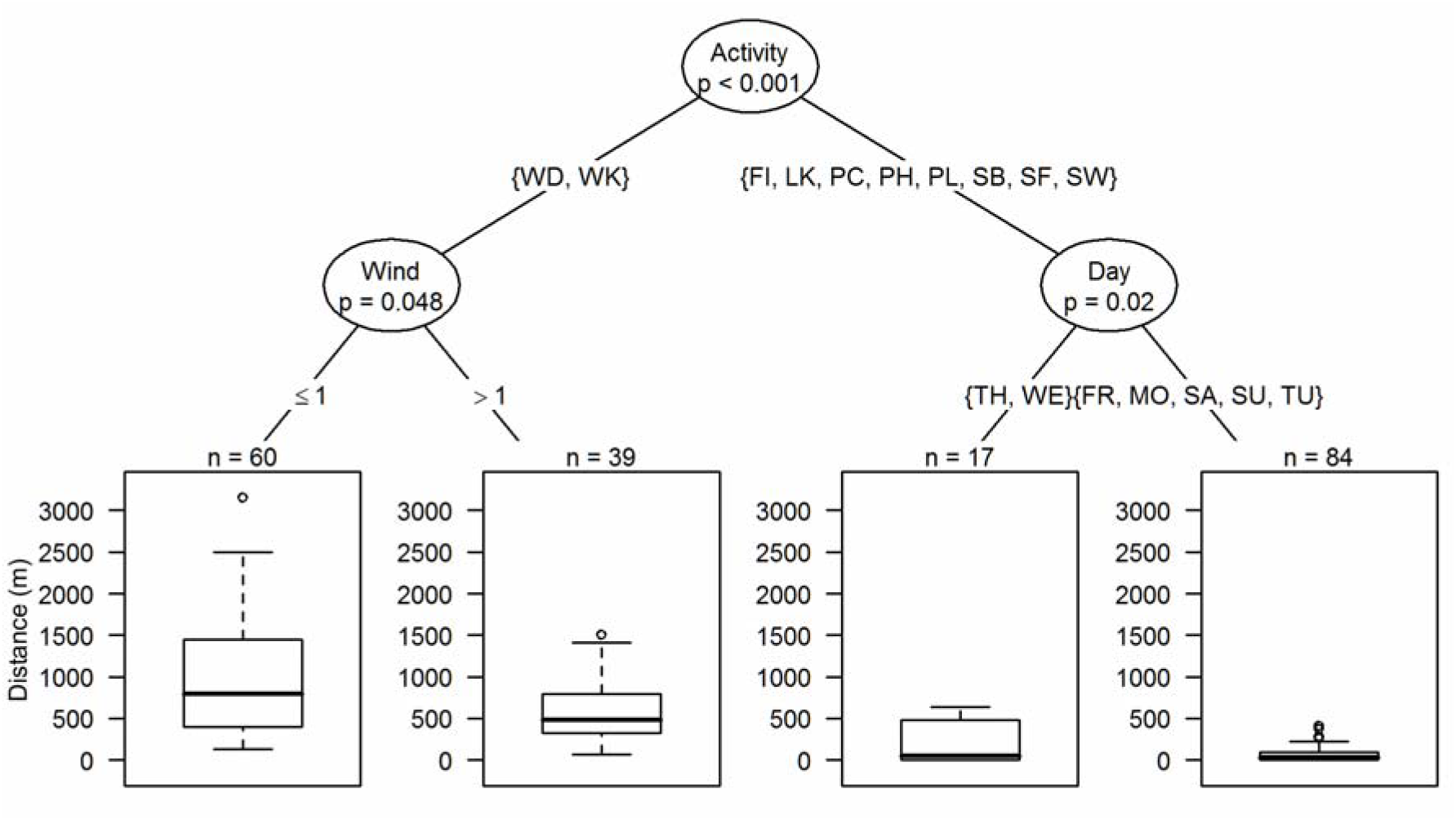
Conditional inference regression tree for distances people walked. The left branch indicates that:1) beach walkers (WD = dog walking, WK = without dogs) typically covered longer distances than did other beach users (right branch), and 2) distances walked decreased at higher wind speeds (> 1 = light to moderate; stronger winds were not observed).,

The regression tree model resulted in a 38% reduction in cross-validated deviance relative to the mean distance walked null model, however the large 443 m residual RMSE indicates large among person/group variability in distances walked.

## DISCUSSION

This study purposively selected a variety of beaches and sites and, although sampling was somewhat haphazard, the dataset was fairly well-balanced among urban and non-urban sites, weekdays and weekends, high and low tides and gender of beach users. It is suggested that the sample was reasonably representative of morning, mostly local beach users in late summer in northern NSW. It would be easy to conduct similar studies for other regions and at other times of the year, if necessary.

Walking, with or without dogs, was the most frequent beach recreation activity observed in this study. Walking has also been reported to be the most frequent activity for other beaches and coastal wetlands in Australia (Dowling & Weston 1999, Antos *et al.* 2007, Glover *et al.* 2011; Maguire *et al.* 2011) although driving can take over on beaches where 4WDs are permitted (Schlacher *et al.* 2013).

The 95th percentile distance for beach walkers and dogs walkers combined was 1990 m. Applying this result, it is recommended that beach access points be approximately two km distant from shorebird habitat to reduce human intrusions into such zones. Of course, the beach should be longer than two km and less-developed for this separation distance to be implemented. Such beaches are also where larger numbers of beach nesting birds can be found (*e.g.* the Australian Pied Oystercatcher Key Management Sites that were noted in the Introduction).

Compared to flight initiation distances, a two km separation distance seems extremely large but is consistent with the ecology of beach nesting birds and is supported by anecdotal evidence. In particular, beach nesting Australian pied oystercatchers occupy linear territories that can be hundreds of metres in length. For example, the maximum recorded oystercatcher breeding density for South Ballina Beach was 18 pairs in 2000 (NSW Department of Planning, Industry and Environment, *unpubl. data*), over 18.7 km of available habitat, which is equivalent to a mean territory length of 1040 m. Large separation distances can protect such long territories from human intrusions.

Oystercatcher nest location data from 2001–2017 show a 4.3 km gap in the distribution of breeding oystercatchers around the village of Patch’s Beach (currently with 19 dwellings) on South Ballina Beach (NSW Department of Planning, Industry and Environment, *unpubl. data*). Harrison (2009) suggested that this gap was the result of permanent and daily human recreation disturbance which deters oystercatchers from occupying otherwise suitable habitat close to Patch’s Beach. The equivalent 2150 m disturbance radius (= 0.5 × 4300) is close to the 1990 m 95th percentile distance from this study.

Finally, it is suggested that human-wildlife coexistence strategies are a “last-resort” option, for overdeveloped beaches that have already been let down by poor land-use planning and where human recreation is established and prevalent. While coexistence strategies could deliver positive conservation outcomes, managing human behaviour is known to be difficult and costly. Moreover, there is a danger that planners will “cherry pick” elements of coexistence strategies to further the development and recreational use of beaches.

## ACKNOWLEDGEMENTS

This manuscript was submitted to *Stilt* but finally withdrawn. The one reviewer was generally supportive of this work. The Editor was too demanding.

